# Automation for the identification of *Pseudomonas aeruginosa:* Comparison of TDR-300B, VITEK^®^2 and VITEK^®^-MS

**DOI:** 10.1101/510107

**Authors:** Lucky H Moehario, Hans P Boestami, Daniel Edbert, Enty Tjoa, T Robertus

**Author notes:** Correspondence should be addressed to Lucky H Moehario (LHM). These authors contributed equally to this work (Lucky H Moehario, Hans P Boestami, Daniel Edbert). These authors also contributed equally to this work (Enty Tjoa, T Robertus).

## Abstract

Quick and precise methods have always been needed in the medical field to correctly identify the agent of infection. Automated systems for diagnosis of infectious pathogen such *Pseudomonas aeruginosa* from critical patients with infections in the Intensive Care Unit is essential. This study aimed to compare the capability of automated biochemistry-based identification system, TDR-300B and VITEK^®^2, to the one of Matrix-Assisted Laser Desorption/Ionization - Time of Flight (MALDI-TOF), VITEK^®^-MS, in the identification of *P. aeruginosa*.

Samples were *P. aeruginosa* isolates collection from Laboratory of Microbiology, Faculty of Medicine and Health Science UNIKA Atma Jaya. These isolates were refreshed; one single colony of oxidase-positive Gram-negative rods was further inoculated in TDR-300B NF-64 cards and VITEK^®^2 GN cassettes. The bacterial identification was also carried out using VITEK^®^-MS as gold-standard. Positivity of TDR-300B and VITEK^®^2 in the identification of *P. aeruginosa* was 87.09% (27/31) and 90.32% (28/31) in the species level, and 87.09%, 96.77% in the genus level respectively. The congruity of TDR-300B/ VITEK^®^2 in the species and genus level was 83.87% and 87.09%. When compared to VITEK^®^-MS, congruence of VITEK^®^2 was 93.30% (24/26) and TDR-300B was 80.76% (21/26). Sensitivity value for TDR-300B and VITEK^®^2 was high, 95.45% and 100%, positive predictive value and accuracy were lower in TDR-300B than VITEK^®^2; Fisher’ exact value was >0.05, thus there were no significant differences in the capability of TDR-300B and VITEK^®^2 in the identification of *P. aeruginosa*.

## Introduction

Microbiology diagnostic has developed and reached to a state that most microbiology work up algorithms are automated and computerised. The automation of laboratory work has made the time to result reduced and the quality of the results improved [1, 2, 3]. Approximately 70% of medical decision dependent on diagnostics test/laboratory results [4, 5]. Significant increase of sample volume yet limited budgets and personnel shortages forced most laboratories to optimise their workflow, productivity, and analytical quality. At the beginning of automation era, it was not considered to be applicable in the microbiology area of works since too many variables exist, including the complexity and variability of sample types, many different analytical processes and varying volume of samples. Many steps involved in the medical microbiology investigation, starting from inoculation of the specimen, incubation, observing and picking colony up to the identification, and susceptibility testing. Thus, to put this into perspective, there is a need to govern each step performed.

Advance technology breakthrough eases the laboratory procedures; VITEK^®^2 (BioMerieux™) and TDR-300B (Mindray Medical International Limited) are among the automation systems developed for bacteriology diagnostic. Many investigations have been conducted on VITEK^®^2 and, they showed the system was to give reliable results [6–9]. On the other hand, despite having similar system that is colorimetry based on the biochemical test array TDR-300B is relatively new and its performance has not been well reported [10]; one study thus far was reported by Sugiartha et al, 2017 [11]. In the developing country like Indonesia, this colorimetry-based identification system seems appropriate. Other approaches in the identification of microorganisms i.e. matrix assisted laser desorption ionization - time of flight mass spectrometry (MALDI-TOF MS) has emerged as a potential tool for microbial identification and diagnosis which can cover a wide spectrum of examination related to microorganisms ranging from the identification of species and strain to antibiotic resistance, epidemiology studies, biological warfare agents and many more [12]; VITEK^®^-MS is one of the systems.

*Pseudomonas aeruginosa* (*P. aeruginosa*) is an aerobic Gram negative motile bacillus that has been associated for Hospital acquired infection (HAI), especially the one related with the use of ventilator in the Intensive care unit (ICU) [13–15]. A report by Radji et al, 2011 in one hospital in Jakarta, *P. aeruginosa* was the most frequent microorganisms (26%) isolated from respiratory tract infection in ICU [13]; in Banda Aceh, 20% of ventilator associated pneumonia was due to *P. aeruginosa* [14]. Ventilator associated pneumonia (VAP) developed in 28% of the patient receiving mechanical ventilation, and high mortality rate of VAP was due to *P. aeruginosa* multi drug resistant (MDR) strains [15–18]. The presence of such bacteria urges the laboratory to bring forth the cause of infection rapidly and accurately. In the present study we compared the capability of TDR-300B and VITEK^®^2 to VITEK^®^-MS in the identification of *P. aeruginosa*.

## Materials and methods

The study was an analytical study to correlate identification capabilities of TDR-300B and VITEK^®^2 to VITEK-MS, carried out in the laboratory of microbiology of Faculty of Medicine and Health Science, Atma Jaya Catholic University of Indonesia. Ethical approval number: 11/05/KEP-FKUAJ/2017.

### Samples

The samples were *P. aeruginosa* isolates collection stored at -20^°^C in the microbiology laboratory of the UNIKA Atma Jaya since 2015 up to 2018. The isolates were originally isolated from clinical specimens and had been subjected to the conventional diagnostic procedures and identified.

### Microbiology Work Up

The isolates were subcultured on blood agar and MacConkey. After 18-24 hours of incubation at 35°C, a single colony was picked, subcultured into nutrient agar slant, and Gram stained. Colonies picked were Gram negative bacilli, MacConkey negative, oxidase positive.

#### Bacterial Identification Using TDR-300B (NF-64 Card)

Colonies on nutrient agar slant were made into suspension of 0.5 McFarland. Biochemical test array of NF-64 card (18 tests) was shown in Table 1. The inoculated card was then incubated at 37°C for 16-24 hours. Microorganism Analysis System version 1.0.0.7 software was used for reading.

#### Bacterial Identification using VITEK^®^2 (GN cassette)

Bacteria suspension of 0.5 to 0.63 McFarland was prepared for the identification. Biochemical test array (47 tests) was shown in Table 1. The analysis was performed using VITEK^®^2 software system 07.01.

#### Bacterial Identification using VITEK-MS

Pure isolates on nutrient agar was prepared. A direct cell profiling for Gram negative bacteria was performed, in which a single colony of microorganism is picked and spotted directly on to the sample plate and immediately overlaid with the matrix solution. The MALDI-TOF MS used was VITEK^®^-MS (BioMerieux™).

### Analysis

Congruence between TDR 300B and VITEK^®^2 was analysed using Interrater Reliability test. Results expected were Percent of agreement (Pa) or Kappa value. In comparison to VITEK^®^-MS as gold standard, Fisher’s exact test was used and Bayes’ formulas were applied.

**Table 1.** Biochemical tests array of TDR-300B card NF-64 and VITEK^®^ 2 GN cassette.

| **TDR-300B card**<br>**NF-64 (18 tests)** | **VITEK®2 GN cassette (47 tests)** |  |  |
| --- | --- | --- | --- |
| Arginine (ADH) | Ala-Phe-Pro-Arylamidase | β-Xylosidase (BXYL) | Succinate alkaniza- |
| Galactose (GAL) | Adonitol (ADO) | β-Alanine arylami-<br>dasepNA (BALap) | β-N-Acetyl-Galac-<br>tosaminidase<br>(NAGA) |
| Gelatin (GEL) | L-Pyrrolydonyl-Arylami-<br>dase (PyrA) | L-ProlineArylamidase<br>(ProA) | α-galactosidase<br>(AGAL) |
| Xylose (XYL) | L-arbitol (IARL) | Lipase (LIP) | Phosphate (PHOS) |
| Maltose (MAL) | D-cellobiose (dCEL) | Palatinose (PLE) | Glycine arylamidase<br>(GlyA) |
| Acetamide (ACM) | β- Galactosidase (BGAL) | Tyrosine Arylamidase<br>(TyrA) | Ornithine decar-<br>boxylase (ODC) |
| Lysine (LDC) | H2S production (H2S) | Urease (URE) | Lysine decarboxy-<br>lase (LDC) |
| Sodium Citrate (CIT) | β-N-Acetyl-Glu-<br>cosaminidase (BNAG) | D-Sorbitol (dSOR) | L-histidine assmila-<br>tion (IHISa) |
| 2-Nitrophenyl β-D-galactopyranoside<br>(BGA) | GlutamylArylamidase-Transferase (AGLTp) | Saccharose/Sucrose<br>(SAC) | Coumerate (CMT) |
| Lactose (LAC) | D-glucose (dGLU) | D-Tagatose (sTAG) | β-glucuronidase<br>(BGUR) |
| Mannitol (MAN) | γ-Gutamyl- Transferase<br>(GGT) | D-Trehalose (dTRE) | 0/129 resistance<br>(comp.vibrio)<br>(0129R) |
| Glucose (GLUF) | Fermentation Glucose<br>(OFF) | Sodium Citrate (CIT) | Glu-Gly-Arg-Ary-<br>lamidase (GGAA) |
| Urea (URE) | β-Glucosidase (BGLU) | Malonate (MNT) | L-Malate assmila-<br>tion (IMILTa) |
| Esculin (ESC) | D-maltose (dMAL) | 5-Keto-D-Gluconate<br>(5KG) | Ellman (ELLM) |
| Glucose (GLU) | D-mannitol (dMAN) | L-Lactate alkaniza-<br>tion (ILATk) | L-Lactate assmila-<br>tion (ILATa) |
| Sucrose (SUC) | D-mannose (dMNE) | α-glucosidase (AGLU) | |
| Fructose (FRU) |  |  |  |
| Sodium Chloride<br>(NaCl) |  |  |  |

## Results

A total of 52 clinical isolates collection labelled as *P. aeruginosa* was retrieved. After subculture only 30 isolated can be further processed. For positive control, *P. aeruginosa* ATCC 27853 was used. Positivity of TDR-300B in the identification of *P. aeruginosa* was 87.09% (27/31) 27 positive results included ATCC 27853 (Table 2);. The average probability of the phenotype based on the 18 biochemical tests shown was 96.14% (CI 95%; SD 8.1%). Of these 27 isolates identified as *P. aeruginosa*, there was 4 isolates showed probability value below deviation range, and produced different biochemical results toward the following: galactose, 2-nitrophenyl beta-D-galactopyranoside, mannitol, glucose and fructose. Further, the TDR-300B misidentified 4 isolates, they were identified as *Shewanella putrefaciens, Chryseobacterium meningosepticum, Stenotrophomonas maltophilia and Acinetobacterbaumanii* (Table 2).

Positivity of VITEK^®^2 in the species level was 90.32% (28/31) (Table 2); mean probability was 96.97% (CI 95%; SD 3.98%). Of those identified as *P. aeruginosa*, 1 isolate showed probability value lower than deviation range. Differences in the biochemical reactions occurred to the following tests: beta-N-acetyl-glucosaminidase, D-mannitol, lipase, urease, D-trehalose, coumerate, O/129 resistance (comp. vibrio), Glu-Gly-Arg-arylamidase, and L-lactate assimilation.

**Table 2.** Identifications of *P. aeruginosa* by TDR-300B, VITEK^®^2, and VITEK^®^-MS.

| No. | TDR-300B |  | VITEK®2 |  | VITEK®-MS |
| --- | --- | --- | --- | --- | --- |
|  | Identification | Probability<sup>a</sup> | Identification | Probability | MS |
| ATCC 27853 | *P. aeruginosa* | 99,72% | *P. aeruginosa* | 96% | *P. aeruginosa* |
| 1 | *P. aeruginosa* | 99,25% | *P. aeruginosa* | 95% | *P. aeruginosa* |
| 2 | *S. putrefaciens* | 75,85% | *P. aeruginosa* | 87% | *P. aeruginosa* |
| 3 | *P. aeruginosa* | 99,72% | *B. pseudomallei* | 91% | *P. aeruginosa* |
| 4 | *P. aeruginosa* | 92,94% | *P. aeruginosa* | 89% | *P. aeruginosa* |
| 5 | *P. aeruginosa* | 83,50% | *P. aeruginosa* | 99% | *P. aeruginosa* |
| 6 | *P. aeruginosa* | 99,57% | *P. aeruginosa* | 99% | *P. aeruginosa* |
| 7 | *P. aeruginosa* | 83,50% | *P. aeruginosa* | 97% | *P. aeruginosa* |
| 8 | *P. aeruginosa* | 99,25% | *P. aeruginosa* | 91% | *P. aeruginosa* |
| 9 | *P. aeruginosa* | 66,36% | *P. putida* | 95% | *P. putida* |
| 10 | *P. aeruginosa* | 99,54% | *P. aeruginosa* | 99% | *P. aeruginosa* |
| 11 | *P. aeruginosa* | 99,54% | *P. aeruginosa* | 90% | *P. aeruginosa* |
| 12 | *C. meningosepticum* | 58,27% | *P. aeruginosa* | 93% | *P. aeruginosa* |
| 13 | *P. aeruginosa* | 99,72% | *P. aeruginosa* | 99% | *P. aeruginosa* |
| 14 | *P. aeruginosa* | 99,36% | *P. aeruginosa* | 99% | *P. aeruginosa* |
| 15 | *P. aeruginosa* | 99,84% | *P. aeruginosa* | 99% | *P. aeruginosa* |
| 16 | *P. aeruginosa* | 99,74% | *P. aeruginosa* | 99% | *P. aeruginosa* |
| 17 | *S. maltophilia* | 99,95% | *P. aeruginosa* | 99% | *P. aeruginosa* |
| 18 | *P. aeruginosa* | 99,25% | *P. aeruginosa* | 99% | *P. aeruginosa* |
| 19 | *A.baumannii* | 70,54% | *P. putida* | 95% | *P. aeruginosa* |
| 20 | *P. aeruginosa* | 99,54% | *P. aeruginosa* | 99% | *P. aeruginosa* |
| 21 | *P. aeruginosa* | 99,72% | *P. aeruginosa* | 95% | *P. aeruginosa* |
| 22 | *P. aeruginosa* | 82,55% | *P. aeruginosa* | 97% | *P. aeruginosa* |
| 23 | *P. aeruginosa* | 99,72% | *P. aeruginosa* | 98% | Not tested |
| 24 | *P. aeruginosa* | 99,54% | *P. aeruginosa* | 95% | Not tested |
| 25 | *P. aeruginosa* | 99,84% | *P. aeruginosa* | 99% | Not tested |
| 26 | *P. aeruginosa* | 99,84% | *P. aeruginosa* | 97% | Not tested |
| 27 | *P. aeruginosa* | 99,74% | *P. aeruginosa* | 86% | *P. aeruginosa* |
| 28 | *P. aeruginosa* | 99,25% | *P. aeruginosa* | 99% | *P. aeruginosa* |
| 29 | *P. aeruginosa* | 99,25% | *P. aeruginosa* | 99% | Not tested |
| 30 | *P. aeruginosa* | 99,25% | *P. aeruginosa* | 97% | *P. aeruginosa* |

<sup>a</sup>Probability (%) represents the capability of either TDR-300B or VITEK<sup>®</sup>2 in identifying the *P. aeruginosa* based on the positivity of biochemical reactions in each card which has been stored in the data base.

Further, 3 out 31 isolates tested were identified as *Burkholderia pseuodomallei* (1 isolate) *and Pseudomonas putida* (2 isolates) (Table 2). The positivity of both systems in the identification in the genus level was examined; TDR-300B was 87.09% (27/31) and VITEK^®^2 showed 96.77% (30/31).

The biochemical tests array in TDR-300B (NF-64 card) and VITEK^®^2 (GN cassette) has only 9 biochemical tests which are similar i.e. maltose, lysine, sodium citrate, glucose oxidation, glucose fermentation, sucrose/saccharose, urea, mannitol, and xylose. Most of the tests showed either positive or negative results, therefore only Percent of agreement (Pa) can be evaluated. In present study, both cards showed the same negative results for maltose, lysine, sodium citrate, glucose oxidation, sucrose and urea; it showed good to almost perfect agreement (Pa >60%) for these tests. While for mannitol, glucose fermentation, and xylose the agreement was none to fair agreement (Pa 0-40%); The TDR-300B NF-64 card test for mannitol, glucose fermentation, and xylose showed more positive results compared to VITEK^®^2 GN card, therefore resulted in low agreement (Table 3).

The congruity of TDR-300B and VITEK^®^2 was analysed and the results showed the congruence in the species level was 83.87% (26/31); both systems identified 25 isolates identified as *P. aeruginosa*, and both misidentified 1 isolate as *Acinetobacter baumannii* by TDR-300B, and *Pseudomonas putida by* VITEK^®^2; these systems showed incongruence in the identification of 5 isolates in the species level. In the genus level, congruity between the two systems was 87.09% (27/31), in which 26 Pseudomonas genus was identified correctly by both systems and 1 genus was identified as other than Pseudomonas.

**Table 3.** Analysis of agreement from 9 biochemical tests of TDR-300 B and VITEK^®^2.

|  | Same Result | TDR-300 B (+)<br>VITEK®2 (-) | TDR-300 B (-)<br>VITEK®2 (+) | *Pa |  |
| --- | --- | --- | --- | --- | --- |
|  |  |  |  | N | % |
| Xylose | 2 | 29 | 0 | 31 | 6.5% |
| Glucose Fermentation | 9 | 22 | 0 | 31 | 29.0% |
| Mannitol | 12 | 14 | 2 | 28 | 42.9% |
| Urea | 22 | 0 | 9 | 31 | 71.0% |
| Glucose | 28 | 0 | 3 | 31 | 90.3% |
| Maltose | 29 | 0 | 2 | 31 | 93.5% |
| Na Citrate | 29 | 0 | 2 | 31 | 93.5% |
| Sucrose | 29 | 1 | 1 | 31 | 93.5% |
| Lysine | 30 | 1 | 0 | 31 | 96.8% |

\*Percent of agreement

VITEK^®^-MS was also employed for the identification on 26 isolates (Table 2). Out of 26 isolates tested, 25 (96.15%) isolates were identified as *P. aeruginosa*, and 1 (3.84%) isolate as *P. putida;* in the genus level positivity was 100% (26/26). When compared to VITEK^®^-MS, VITEK^®^2 showed the congruence of 93.30% (24/26). The incongruence was shown from 2 isolate, it was identified as *B. pseudomallei*, and *P.putida* by VITEK^®^2 but both were *P. aeruginosa* by VITEK-MS; the TDR-300B showed congruity of 80.76% (21/26), and another 5 isolates were identified as other then *P. aeruginosa* by TDR-300B but all were *P. aeruginosa* by VITEK^®^-MS.

Using the VITEK^®^-MS as gold standard, sensitivity value of TDR-300B was high, 95.45%, and VITEK^®^2 was 100%. The positive predictive value (PPV) and accuracy, however, were lower in TDR-300B than VITEK^®^2 (Table 4). The Fisher’ exact value for both comparison was >0.05, so there was no significant differences in the capability of TDR-300B and VITEK^®^2 in the identification of *P. aeruginosa*.

**Table 4.** Diagnostic capability of TDR-300B and VITEK^®^2 as compared to VITEK^®^ -MS.

|  | **TDR 300B vs VITEK®-MS** | **VITEK 2 vs VITEK®-MS** |
| --- | --- | --- |
| Sensitivity | 95.45%<br>(CI 95%; 77.16% - 99.88%) | 100%<br>(CI 95%; 85.18% - 100.00%) |
| PPV | 84.00%<br>(CI 95%; 82.74% - 85.19%) | 92.00%<br>(CI 95%; 83.78% - 96.24%) |
| Accuracy | 80.77%<br>(CI 95%; 60.65% - 93.45%) | 92.31%<br>(CI 95%; 74.87% - 99.05%) |
| Fischer's exact Value | 1 ($\alpha=0.05$) | 0.1154 ($\alpha=0.05$) |

## Discussion

Pseudomonas is clinically important bacteria and frequently isolated from clinical specimens especially those collected from patients in the ICUs. In the present study, the congruity of TDR-300B and VITEK^®^2 was quite high, both in species and genus level (83.87% and 87.09%); the incongruence was 16.2% derived from 5 isolates as shown in Ta- ble 2. In terms of identification, both TDR-300B and VITEK^®^2 showed an almost similar probability to identify *P. aeruginosa* i.e. 96.14% versus 96.97%. Species tested showed biochemical tests pattern with high confidence above 95%; databases and biochemical contents in each card contribute to the results in the identification of bacteria species. The TDR-300B NF-64 card with only 18 biochemical tests seemed to have more tendency to misidentify *P. aeruginosa* compared to VITEK^®^2 with the GN-cassette which contains 47 biochemical tests. Moreover, our study identified only 9 biochemical tests were similar between the TDR-300B and VITEK^®^2, and 6 of those (i.e. maltose, lysine, sodium citrate, glucose oxidation, sucrose and urea) showed good to perfect agreement, while the other 3 tests (mannitol, glucose fermentation, and xylose) showed low agreement.

In 2014, da Silva Paim et al [8] investigated the performance of VITEK^®^2 and compared with conventional method in the identification of Gram positive cocci. The results showed that VITEK^®^2 correctly identified the species and genus levels at 77.9% and 97.1% respectively of the Staphylococcus sp., Enterococcus. sp. and Streptococcus sp. The report seemed similar to the one conducted in 2002 by Ligozzi et al, in which compared to API Staph and API 20 Strep, the VITEK^®^2 system correctly identified up to the species level, >90% for *S. aureus, S. agalactiae, S. pneumoniae, Enterococcus faecalis, Staphylococcus haemolyticus*, except for *Staphylococcus epidermidis and was* least able to identify *Enterococcus faecium i.e*. 71.4% [6].

Further, the ability of the automated system in the identification Gram negative microorganisms were also carried out. Darbandi, 2010 reported a complete coalition of genus and species levels from 90 clinical isolates (77.8%) identified by API 20E/API 20NE Microsystems and VITEK^®^2 [19]. In 2016 a study conducted by Nayeem-u-din Wani et al, showed overall concordance of 91.1% of manual conventional method versus VITEK^®^2 in Gram negative bacteria identification between [20]; the concordant identification was seen with the isolates of *Klebsiella pneumoniae, Acinetobacter l·woffii, Citrobacter freundii, Enterobacter cloacae, P. aeruginosa* and *Salmonella* Typhi. Hernandez-Duran et al, 2017 compared the usefulness of VITEK^®^2 against the MicroScanWalkAway^®^SI for bacterial identification [9]. They found of the 54 bacterial strains composed of 20 Gram-positive cocci, 34 Gram-negative rods isolated from hospitalised patients and 13 reference strains, 89.5% were successfully identified at the species level by both systems. Capability of TDR-300B and VITEK^®^2 in bacterial identification was studied by Sugiartha et al, in 2017. They reported the accuracy of TDR-300B was 90.9% versus VITEK^®^2 from 33 Gram-negative and Gram-positive bacteria species [11]. In the present study, comparison of TDR-300B and VITEK^®^2 to VITEK^®^-MS as gold standard showed that VITEK^®^2 has higher accuracy (92.31%) and PPV (92.00%) than TDR-300B (80.77% and 84.00%). The sensitivity of TDR-300B and VITEK^®^2 was high, 95.45% and 100% respectively. How-ever, the PPV and accuracy values seemed lower in TDR-300B than VITEK^®^2. Specificity and negative predictive value (NPV) were not calculated since in the present study only isolates previously identified as *P. aeruginosa* were included. Fisher’s exact value showed no significance for TDR-300B and VITEK^®^2 because of the small negative numbers. no significant differences can be deducted.

## Conclusions

TDR-300B and VITEK^®^2 showed quite high congruence in the species and genus level in the identification of *P. aeruginosa*. The VITEK^®^2/VITEK-MS showed congruence of 93.30%, and TDR-300B/VITEK-MS 80.76%. The sensitivity for TDR-300B and VITEK^®^2 was high, 95.45% and 100%. Positive predictive value and accuracy, however, were lower in TDR-300B than VITEK^®^2. There was no significance differences in the capability of TDR-300B and VITEK^®^2 in the identification of *P. aeruginosa*. It is worth noted that there was some differences in species identified by TDR-300B and VITEK^®^2, which might affect the management therapy of patients. Therefore, it is important to conduct good practice of microbiology work up, and minimise improper techniques; addition of more biochemical tests may be useful or use other approaches to confirm the identity of the microorganisms.

## Acknowledgement

We thanked dr. Lely Saptawati, Clinical microbiologist from Public Hospital Dr. Moewardi for the support in laboratory work with VITEK-MS MALDI-TOF.

